# Clinical Viability of Magnetic Bead Implants in Muscle

**DOI:** 10.1101/2022.08.01.502296

**Authors:** Cameron R. Taylor, William H. Clark, Ellen G. Clarrissimeaux, Seong Ho Yeon, Matthew J. Carty, Stuart R. Lipsitz, Roderick T. Bronson, Thomas J. Roberts, Hugh M. Herr

## Abstract

Human movement is accomplished through muscle contraction, yet there does not exist a portable system capable of monitoring muscle length changes in real time. To address this limitation, we previously introduced magnetomicrometry, a minimally-invasive tracking technique comprising two implanted magnetic beads in muscle and a magnetic field sensor array positioned on the body’s surface adjacent the implanted beads. The implant system comprises a pair of spherical magnetic beads, each with a first coating of nickel-copper-nickel and an outer coating of Parylene C. In parallel work, we demonstrate submillimeter accuracy of magnetic bead tracking for muscle contractions in an untethered freely-roaming avian model. Here, we address the clinical viability of magnetomicrometry. Using a specialized device to insert magnetic beads into muscle in avian and lagomorph models, we collect data to assess gait metrics, bead migration, and bead biocompatibility. For these animal models, we find no gait differences post- versus pre-implantation, and bead migration towards one another within muscle does not occur for initial bead separation distances greater than 3 cm. Further, using extensive biocompatibility testing, the implants are shown to be non-irritant, non-cytotoxic, non-allergenic, and non-irritating. Our cumulative results lend support for the viability of these magnetic bead implants for implantation in human muscle. We thus anticipate their imminent use in human-machine interfaces, such as in control of prostheses, exoskeletons, and in closed-loop neuroprosthetics to aid recovery from neurological disorders.

## Introduction

The field of human bionics seeks to advance methods for the high-fidelity control of wearable robots to restore and augment human physicality. Delivering on this objective requires new interfacing strategies for determining intent. Today’s standard interface for extrinsic control of wearable robots is surface electromyography (sEMG). This technique senses myoelectric signals via a skin surface electrode, but suffers from signal noise and drift caused by motion artifacts and impedance variations (Calado et al. 2019; Clancy et al. 2002). While significant advances have been made in robotic control via sEMG (Farina et al. 2017), even myoelectric signals measured using invasive strategies (Weir et al. 2008) cannot fully communicate intended muscle forces or joint movements without also incorporating measurements of muscle length and speed (Zajac 1989). However, a real-time sensing technology has not yet been developed that can reliably measure muscle length and speed in humans.

To address this need, we recently developed a strategy for tracking muscle tissue lengths. This technique, known as magnetomicrometry (MM), uses implanted pairs of magnetic beads to wirelessly track muscle tissue lengths in real time (see Figure 1) (Taylor et al. 2021). In parallel work, we further validate the performance of MM in mobile usage (Taylor et al. 2022), lending additional support to its utility for prosthetic and exoskeletal control. Due to its very low time delay, high accuracy, and minimally invasive nature (Taylor et al. 2019), MM has the potential to provide an improved extrinsic controllability over wearable robots.

**Figure 1:**
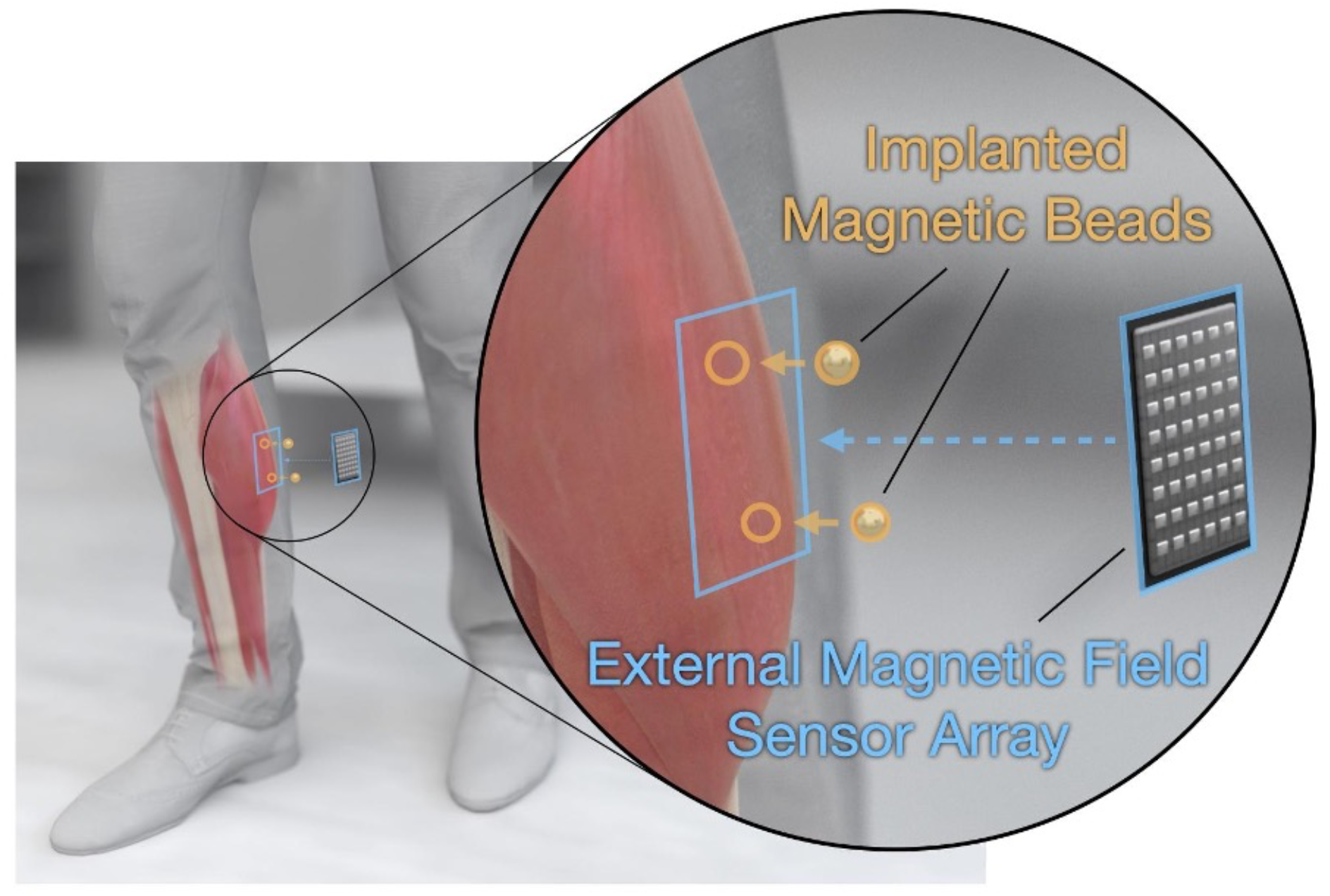
Tracking Muscle Tissue Lengths via Muscle Magnetomicrometry. Magnetomicrometry tracks muscle tissue lengths in vivo, with potential prosthetic and exoskeletal control applications. A surgeon implants a pair of gold- and Parylene-coated magnetic beads (highlighted in orange) into each muscle of interest. An external magnetic field sensor array (highlighted in blue) placed over each pair of implants then senses the passive magnetic fields from the implants, and a computer uses these magnetic field measurements to calculate the positions of the magnetic beads. The computer then calculates the distance between the two beads, providing a real-time estimate of tissue length.

Towards translating MM for human use, we here develop an improved coating strategy for 3-mm-diameter spherical magnetic bead implants and a translatable surgical protocol for implantation. With this new strategy, we apply a first coat of gold and a top coat of Parylene to the magnetic beads, following stringent medical device standards for coating, cleaning, and sterilization. We then address three features that are essential for the clinical use of magnetic bead implants: (1) lack of implant discomfort (2) lack of implant migration, and (3) implant biocompatibility.

### Comfort

These implants are intended for permanent implantation like the tantalum beads used in radiostereographic analysis (RSA). Since the standardization of RSA in 1974 (Selvik 1974), tens of thousands of spherical tantalum beads have been implanted into patients with no adverse reactions (Kärrholm 1989), including into muscle tissues (Clites et al. 2018). However, though the magnetic beads we present here are spherical like those used for RSA, they have a diameter three times larger than RSA beads, necessitating an analysis of implant comfort.

Changes in movement patterns have been routinely analyzed to capture the secondary musculoskeletal effects of discomfort or pain, commonly referred to as antalgic gait (Vincent et al. 2013; Auerbach and Tadi 2021; Nonnekes et al. 2020). Following this previous literature, this work analyzes changes in the percentage of stance phase relative to swing phase in an animal model as a measure of comfort after implantation of these magnetic beads. Specifically, we hypothesize that the percentage of stride time in stance is unaffected by the long-term implantation of the magnetic beads. To evaluate this hypothesis, we use an in vivo turkey model to implant magnetic bead pairs in the right gastrocnemius muscle, and we analyze the percentage of stride time in stance for walking and running before and after implantation.

### Migration

To provide a consistent measurement of tissue length and to ensure patient health, the magnetic implants must (1) not move relative to the surrounding tissue, (2) not move relative to one another, and (3) not migrate out of the muscle. Spherical steel beads and cylindrical titanium-encapsulated magnetic beads implanted individually into the tongue muscle in animal models proved to be stable against migration in short-term studies (6-24 days) (Mimche et al. 2016; Sokoloff et al. 2017). In previous work, we also showed that magnetic beads implanted in pairs do not migrate over a long-term study (191 days) if sufficiently separated from one another at the time of implantation (Taylor et al. 2021). Although we only investigated migration of magnets at a single magnetization strength, we used these results to determine a distance threshold for migration. We then presented a theoretical method for adjusting that threshold based on the strength of the magnets. In this investigation, we hypothesize that distance thresholds for magnetic bead pair migration can be predicted from empirical data of different magnet strengths using a simple model of the force between them. To evaluate this hypothesis, we use an in vivo turkey model to implant magnetic bead pairs in the lateral gastrocnemius and tibialis cranialis muscles, and we monitor long-term magnetic bead positions for migration.

### Biocompatibility

The biocompatibility of magnetic bead implants must also be verified before clinical use. In previous work, as a proof of concept, we used non-clinical-grade Parylene-coated magnets (7 μm coating thickness) with a non-clinical-grade insertion process, and we found only minor inflammation (Taylor et al. 2021). Recent work by Iacovacci et al. (Iacovacci et al. 2021) extends those results, showing that Parylene-coated magnets (10 μm coating thickness) are non-cytotoxic, non-pyrogenic, and systemically non-toxic, resistant to wear under repeated muscle-like mechanical compression, resistant to corrosion in a simulated post-surgical inflammatory environment, and non-irritant under 28-day sub-acute toxicity evaluation. In this study, we investigate the biocompatibility of magnetic beads as long-term implants in muscle. We hypothesize that the implantation of magnetic beads with suitable biocompatible coatings does not cause adverse tissue reactions, using a first coating of nickel-copper-nickel (10 to 25 μm coating thickness), a second coating of gold (at least 5 μm coating thickness), and an outer coating of Parylene C (21 μm coating thickness). Using a bead insertion methodology from a clinical-grade device, we examine tissue responses to the implants. Specifically, the coated magnets are evaluated for irritation using a 2-week and 26-week intramuscular implantation, as well as an intracutaneous protocol. Further, a cytotoxic testing protocol is employed using a 72-hour minimal essential media elution extraction methodology, and sensitization testing is conducted using a non-allergenic testing methodology.

## Materials and Methods

We implanted magnetic beads in ten wild turkeys over an eight-month period. All turkey experiments were approved by the Institutional Animal Care and Use Committees (IACUC) at Brown University and the Massachusetts Institute of Technology. We obtained the wild turkeys (Meleagris gallopavo, adult, seven male, three female) from local breeders and cared for them in the Animal Care Facility at Brown University on an ad libitum water and poultry feed diet.

All biocompatibility testing protocols were reviewed and approved by the WuXi AppTec IACUC prior to the initiation of testing. Twelve rabbits and seventeen guinea pigs were used in the biocompatibility testing portion of this work. Albino rabbits (Oryctolagus cuniculus, young adult, four female, eight male) and albino guinea pigs (Cavia porcellus, young adult male) were obtained from Charles River Laboratories and maintained in the WuXi AppTec animal facility according to NIH and AAALAC guidelines on an ad libitum water and certified commercial feed diet.

### Magnetic Bead Implants

We manufactured the magnetic bead implants following medical device standards for coating, cleaning, and sterilization. SM Magnetics manufactured the base for the magnetic bead implants. The base was a 3-mm-diameter sphere composed of sintered neodymium-iron-boron and dysprosium (grade N48SH) and plated with nickel-copper-nickel (see Figure 2). This base material was delivered unmagnetized to Electro-Spec, Inc., who cleaned the spheres and tumble barrel plated them with 5 μm gold in adherence to ASTM B488, Type III, Code A, Class 5. Specialty Coating Systems (SCS) cleaned these gold-plated beads with isopropyl alcohol and deionized water in an ISO-7 clean room and applied adhesion promoter (AdPro) to the gold surface. SCS then coated the beads with 21 μm Parylene C via vapor deposition polymerization.

**Figure 2:**
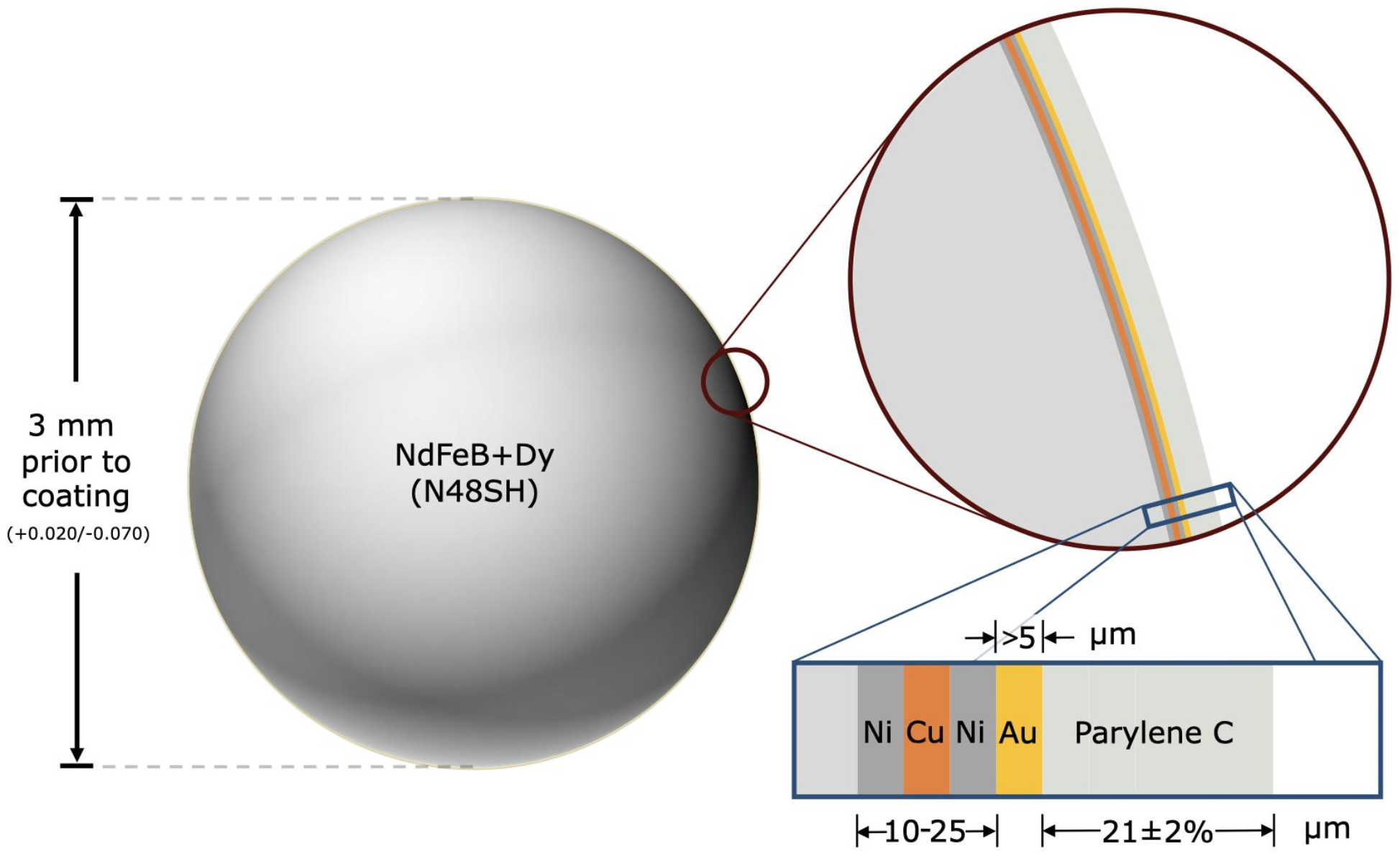
Spherical Magnetic Beads for Implantation in Muscle. The magnetic bead implants are composed of sintered neodymium-iron-boron (Nd:26-33%, Fe:63.2-68.5%, B:1.0-1.2%, Dy-Tb:0-1.5%, Nb:0.3-1.4%, Al:0.1-1.0%), with dysprosium and terbium added to increase the maximum working temperature to 150°C, which allows the implants to be autoclaved when needed. The base of each magnetic bead implant is approximately 3 mm in diameter (surface area ∼0.283 cm^2^) and has an approximate residual flux density of 1.36-1.42 T. The implant is coated in 10 to 25 μm nickel-copper-nickel (see inset with blue border), with the outermost nickel coating being 99.99% pure. It is then coated in at least 5 μm of 99.9% pure gold and approximately 21 μm of Parylene C, resulting in a diameter after coating of approximately 3.001 to 3.129 mm.

To ensure the implants were free from debris, SCS shipped the Parylene-coated beads in double vacuum packaging to KKS Ultraschall AG, who prepared the implants for magnetization by cleaning and packaging them in polyvinyl chloride tubes and caps received from SM Magnetics. Specifically, in their ISO-8 clean room, KKS ultrasonically cleaned these tubes and caps, inserted 32 beads into each tube, placed the caps on each end, and individually double-packaged each tube into two sealed, well-fitting bags. SM Magnetics then placed each double-packaged tube into a magnetizing coil and used a first magnetizing pulse to align the beads, followed by a second magnetizing pulse to magnetize the beads.

To prepare the beads for surgical insertion, Halifax Biomedical, Inc. (HBI) manufactured cartridges to hold the biocompatibly-coated magnetic beads (see Figure 3). For the in-house portion of this work, we placed the beads in these cartridges directly from the tubes via pressing and shearing, then autoclaved the bead set. For outside laboratory biocompatibility testing (WuXi AppTec), HBI inserted the beads into these cartridges (eight beads per cartridge, ISO-7 clean room) directly from the tubes via pressing and shearing, triple-packaged each cartridge into a clamshell container, and sterilized each bead set via gamma sterilization before providing them to the biocompatibility testing facility.

**Figure 3:**
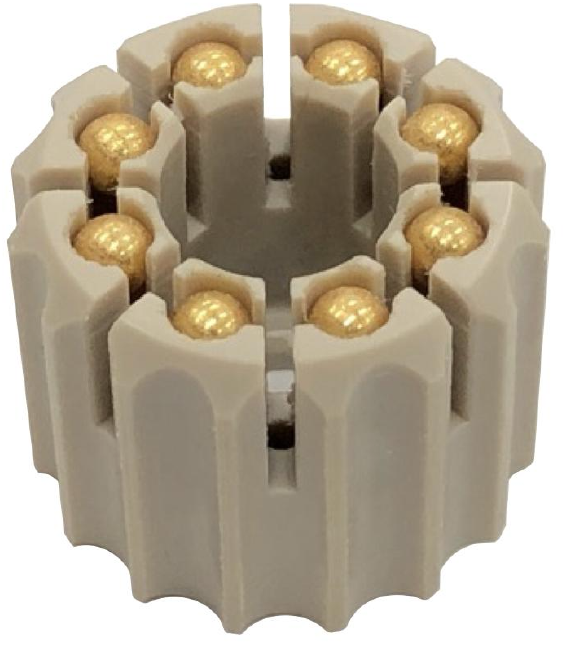
Magnetic Bead Set. Each magnetic bead set consists of eight coated, cleaned, and magnetized beads subsequently inserted into a magnetic bead cartridge for packaging, sterilization, storage, and deployment. The surgeon deploys the magnetic beads directly from the cartridge using the custom Halifax magnetic bead insertion device (see Figure 4).

### Insertion Device

We contracted HBI to manufacture a customized magnetic bead insertion device (see Figure 4) using the same components used in their tantalum bead inserter, but with the shaft and pushrod made from medical-grade titanium alloy to prevent the magnetic beads from sticking to the metal during the implantation procedure. The shaft and pushrod were also widened slightly to accommodate a larger bead compared to standard tantalum bead implants and shortened to simplify the implantation procedure for the magnets, which are not inserted as deeply as tantalum beads. In addition, the 8-bead magnetic bead set was designed to be advanced by two clicks between insertions, allowing it to be a drop-in replacement for the typically-used tantalum bead sets containing 16 smaller beads. For the outside laboratory biocompatibility testing, we provided the biocompatibility testing facility with the insertion device.

**Figure 4:**
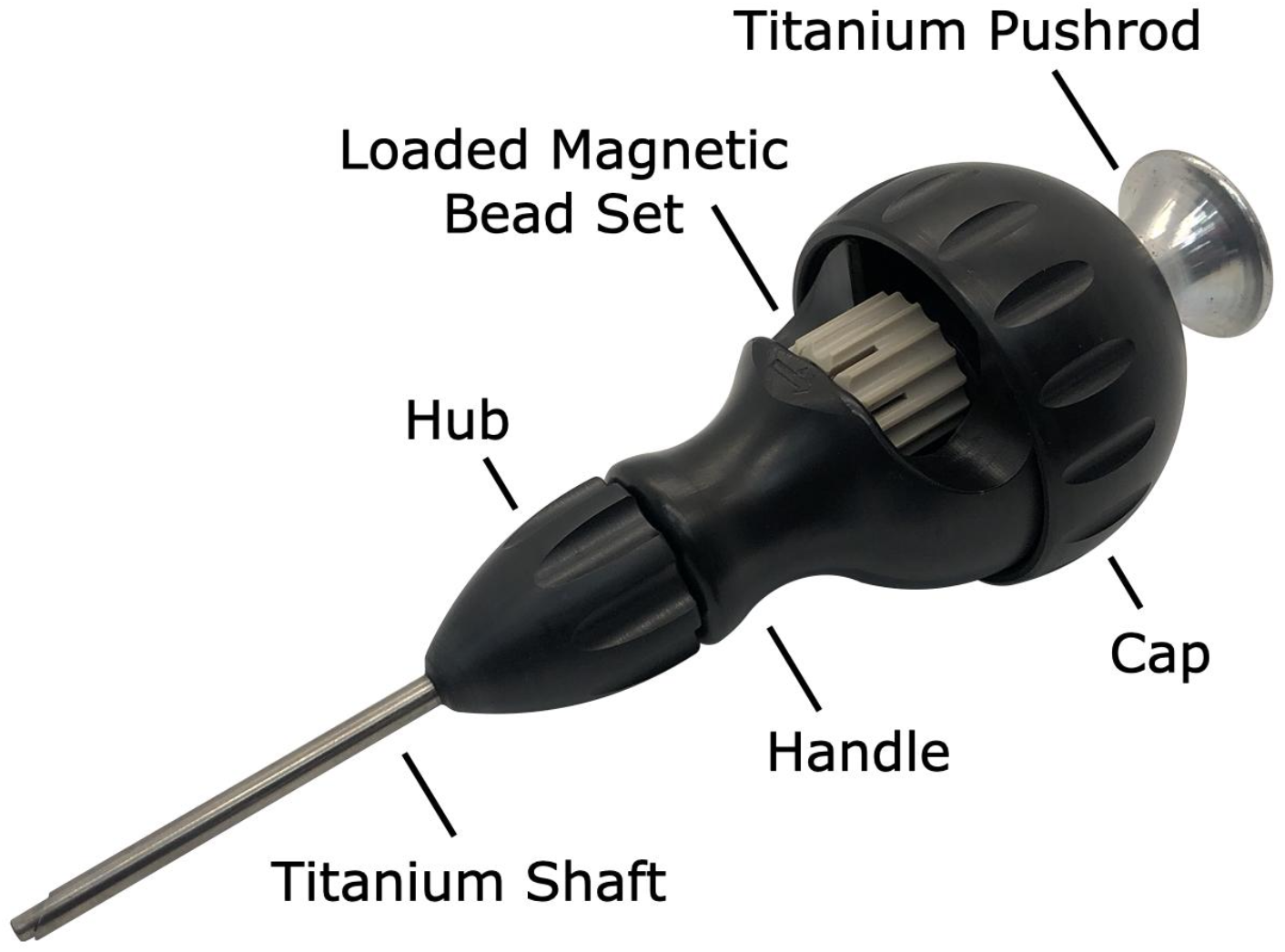
Custom Halifax Magnetic Bead Insertion Device. A custom magnetic bead insertion device enables the surgeon to implant the magnetic beads directly into the muscle without directly handling the implants. The device reuses the design for tantalum bead insertion already clinically in use, widening the shaft and pushrod to accommodate the 3-mm-diameter beads and replacing the shaft and pushrod with titanium to eliminate any attraction of the magnetic beads to the device materials. A replaceable cap allows a new magnetic bead set to be loaded into the device, and an insertable pushrod allows the surgeon to deploy a magnetic bead from the magnetic bead set into the muscle via the shaft. The hub unscrews from the handle to facilitate thorough cleaning and sterilization.

### Implantation

We performed all implantation procedures under sterile conditions in spaces approved for veterinary interventions. For each procedure, we placed the turkey under general anesthesia and draped the turkey in typical sterile fashion, as described in previous work (Taylor et al. 2021). We identified the target muscle via surface anatomic landmarks and further confirmed its location via ultrasound. We then identified specific insertion sites within the muscle, ensuring an intermagnet distance greater than 3 cm, and marked these planned insertion sites with a sterile operative pen. At each insertion site, we made a 1 cm length incision through all layers of skin using a No. 11 surgical scalpel. We then dissected through the underlying subcutaneous fat layer down to the underlying fascia, then incised the fascia with the No. 11 surgical scalpel to expose the underlying target muscle. Finally, we made a 1 cm length incision through the epimysium to expose the underlying muscle fibers.

After loading a magnetic bead set into the Halifax Insertion Device, we marked the planned implantation depth for each insertion site (approximately 6 mm for the proximal site and 2-3 mm for the distal site) on the shaft of the insertion device and on a pair of fine surgical scissors (using a sterile operative pen and sterile ruler). With the aid of the fine surgical scissors, we pretunneled the intramuscular insertion path to the marked depth. We then placed the shaft of the Halifax magnetic bead inserter into the pretunneled path, using the depth marking as a guide. Keeping the insertion device in position, we deployed each magnetic bead implant by firmly grasping the insertion device and inserting the pushrod into the device until the knob of the pushrod contacted the cap, then withdrew the insertion device with the pushrod still in contact with the cap. Additional surgical instruments were utilized as necessary due to animal-specific anatomy (i.e., when the muscle was found to be too thin or too shallow).

After insertion, we reapproximated the epimysial edges at each site using 4-0 chromic interrupted stitches. While suturing, we took care to not bring ferromagnetic surgical instruments closer than necessary to the implants to avoid inadvertent displacement of the magnetic beads. Finally, we confirmed the implantation depth and distance between the magnetic beads via MM and ultrasound, approximated the skin edges at each implantation site using 4-0 polypropylene interrupted percutaneous stitches, then confirmed the magnetic bead positions once again.

### Comfort

We defined comfort as the absence of change in the percentage of stride time in the stance phase for the walking and running animals following magnetic bead implantation. Following the same techniques listed above, we implanted a magnetic bead pair in the mid-belly of the right gastrocnemius of the three female turkeys.

To minimize variability in movement patterns, we trained each turkey to walk and run on an enclosed treadmill at 1.5, 2.0, 2.5, 3.0, and 3.5 m/s. In total, there were 15 training sessions: nine times prior to implantation and six times following implantation. During training, we randomized treadmill speed every two minutes for a total of ten minutes. We collected pre-implantation biomechanics at session 9 and post-implantation biomechanics at session 15 (three weeks post-implantation) on a high-speed camera (Flare 12M180MCX, IO Industries) at 120 Hz. For each speed, we visually confirmed that the turkey’s movement pattern had stabilized before collecting data.

We also manually identified toe strikes and toe offs for at least 20 consecutive gait cycles at each speed, during selected periods of gait that visually minimized irregular movements (e.g., jumping, standing).

Post process, we confirmed consistent movement patterns by excluding strides that were greater than three standard deviations from the mean percentage of right leg stride time in stance, then subsequently replaced inconsistent strides with the next stride recorded.

After collecting the data, we performed statistical tests to determine whether there was an effect of surgery or speed on percentage of stride time in stance using SPSS Statistics 27 (IBM). A linear mixed model tested for the fixed main effect of speed (i.e., 1.5, 2.0, 2.5, 3.0, and 3.5 m/s) and fixed main effect of unilateral implants (i.e., pre-surgery and post-surgery) on the percentage of right leg stride time in stance. Using the theory of generalized estimating equations (Liang and Zeger 1986; Lipsitz et al. 1994), the linear mixed model for estimating means in a repeated measures model is robust to non-normality of the outcomes, so our estimated effects can be considered unbiased estimates. When significant main effects were found, post-hoc Tukey tests identified pairwise differences between individual speeds. We report effect sizes as *ηp*2 and Cohen’s *d* for main effects and pairwise comparisons, respectively. Values of *d* > 0.2, 0.5, and 0.8 indicate small, moderate, and large effects, respectively (Cohen 2013). All statistical tests used an alpha level of 0.05.

### Migration

In the seven male turkeys, we intentionally placed the magnetic beads at a range of distances and in multiple muscles to explore different circumstances in which the magnetic beads might migrate toward one another. The distance between magnets was measured as described in previous work (Taylor et al. 2021), except that CT scans were performed over a longer time period (immediately following implantation and then again after eight months).

### Biocompatibility

To analyze tissue response to the clinical-grade implants inserted using the Halifax magnetic bead inserter, in-house observations were made first, with a total of eight histological samples taken from four of the male turkeys, each at eight months post-implantation. Histology was performed as in previous work (Taylor et al. 2021), with the addition of Masson’s Trichrome staining to highlight the fibrous tissue.

To test the device biocompatibility under good laboratory practice (GLP) compliance (USFDA, Code of Federal Regulations, Title 21, Part 58 - Good Laboratory Practice for Nonclinical Laboratory Studies), we submitted fully manufactured magnetic bead sets and insertion devices to WuXi AppTec for intramuscular implantation, cytotoxicity, intracutaneous irritation, and sensitization testing. All magnetic beads used in the testing were deployed from the magnetic bead cartridges using the insertion device, and all tests were conducted in compliance with international standard ISO 10993-12:2012, Biological Evaluation of Medical Devices, Part 12: Sample Preparation and Reference Materials.

### Biocompatibility: Intramuscular Implantation Testing

To test for local effects of the magnetic beads contacting skeletal muscle, three magnetic beads and three negative control articles (high density polyethylene, HDPE) were implanted into the left and right paravertebral muscles, respectively, in nine rabbits. Four rabbits were used for the two-week implantation test, and the remaining five rabbits were reserved for the twenty-six-week implantation test. At the test endpoints, the muscles were explanted, fixed in 10% neutral buffered formalin, and sent to the study pathologist for analysis.

Each implantation site was analyzed via clinical, gross, and histopathologic information. Each site was examined microscopically and scored for inflammation and tissue response on a scale from 0 (no inflammation/response) to 4 (severe inflammation/response). The average of the magnetic bead site scores minus the control article site scores was used to determine an irritant ranking score, with a negative value considered 0.0. A score from 0.0 up to 2.9 was considered to indicate that the magnetic bead implants were non-irritant. The intramuscular implantation tests were conducted in accordance with the ISO Test Method for Implantation in Muscle, ISO 10993-6: 2016, Biological Evaluation of Medical Devices - Part 6: Tests For Local Effects After Implantation.

### Biocompatibility: Cytotoxicity Testing

To test for toxicity of the implants to mammalian cells, 54 magnetic beads were extracted at a ratio of 3 cm^2^ / 1 mL (surface area per volume) in Eagle’s minimal essential medium (E-MEM) supplemented with 5% volume-per-volume fetal bovine serum (FBS) for 72 hours at 37°C, then shaken well. A negative control (HDPE) extraction, positive control (0.1% zinc diethyldithiocarbamate, ZDEC, polyurethane film) extraction, and cell control were also prepared in parallel to and under equivalent conditions as the magnetic bead extraction.

Completed extractions were added to fully-formed cell culture wells containing L-929 mouse fibroblast cells (American Type Culture Collection, ATCC # CCL-1) after removal of the culture maintenance medium. Specifically, 1 mL of each extraction was inserted into each of three cell culture wells, and the cell culture wells were incubated in a humidified atmosphere for 72 hours.

The cell cultures were examined microscopically and scored for cytopathic effects (lysis, crenation, plaques, and excessive rounding of cells) at 24, 48, and 72 hours on a scale from 0 (no reactivity) to 4 (severe reactivity). A score of 0, 1, or 2 was considered to indicate that the magnetic bead implants were non-cytotoxic. This cytotoxicity testing was conducted in compliance with ISO 10993-5:2009, Biological Evaluation of Medical Devices, Part 5: Tests for *In Vitro* Cytotoxicity.

### Biocompatibility: Sensitization Testing

To test for the induction of allergic reactions, magnetic beads were extracted at a ratio of 3 cm^2^ / 1 mL (surface area per volume) into each of 5.1, 8.1, and 10.1 mL of 0.9% normal saline (54, 86, and 107 magnetic beads) and 5.1, 8.1, and 10.1 mL of sesame oil (54, 86, and 107 magnetic beads). These extractions were freshly prepared for corresponding phases of the test and were performed over 72 hours at 50°C, with agitation during the course of the extraction. Vehicle controls (containers of liquid without magnetic beads) were prepared in parallel to and under equivalent conditions as the magnetic bead extractions.

One tenth of one milliliter of each extraction was injected intracutaneously into the dorsal dermis of eleven guinea pigs (one on each side of the midline). Adjacent these first two injections, two additional 0.1 mL injections of an immunostimulant (equal parts Freund’s Complete Adjuvant and 0.9% sterile saline) were injected subcutaneously. An additional 0.05 mL of each extraction was then mixed with 0.05 mL of the immunostimulant, and each was injected subcutaneously adjacent the first injections. All six injections were repeated using negative (vehicle) controls in place of the magnetic bead extractions in each of six additional guinea pigs. Six days post-injection, the injection sites were treated with 10% sodium lauryl sulfate in mineral oil. After 24 hours, the sites were then cleaned, and a 2×4 cm filter paper saturated with magnetic bead extract (or vehicle control, for the negative control group) was applied dermally to the site for 48 hours, then removed.

After an additional fifteen days, two 2×2 cm filter papers saturated with magnetic bead extract and vehicle control, respectively, were applied dermally to the right and left flanks, respectively, of each animal for 24 hours, then removed. At 24 to 48 hours after patch removal, these 2×2 cm exposure sites were observed and scored for irritation and sensitization reaction on a scale from 0 (no erythema/edema) to 3 (intense erythema/edema). A score of 0 (or a score not exceeding the most severe negative control reaction, if non-zero) was considered to indicate that the magnetic bead implants did not elicit a sensitization response. This sensitization testing was conducted in compliance with ISO 10993-10: 2010, Standard, Biological Evaluation of Medical Devices, Part 10: Tests for Irritation and Skin Sensitization.

### Biocompatibility: Intracutaneous Irritation Testing

To test for local irritation of dermal tissue due to potential extractables or leachables, magnetic beads were extracted at a ratio of 3 cm^2^ / 1 mL (surface area per volume) into each of 6 mL of 0.9% normal saline (64 magnetic beads) and 6 mL of sesame oil (64 magnetic beads). These extractions were performed over 72 hours at 50°C, with agitation during the course of the extraction, and vehicle controls (containers of liquid without magnetic beads) were prepared in parallel to and under equivalent conditions as the magnetic bead extractions.

One milliliter of each extraction and of each vehicle control was then injected intracutaneously into the dorsal dermis in each of three rabbits. Specifically, the magnetic bead extractions were injected to the right of the midline, and the vehicle controls were injected to the left of the midline. Each extraction was delivered as five 0.2 mL injections at locations spatially distributed along the cranial-caudal axis, with the normal saline extractions being delivered medial to the sesame oil extractions.

The injection sites were observed and scored for gross evidence of erythema and edema at 24, 48, and 72 hours on a scale from 0 (no erythema/edema) to 4 (severe erythema/edema). A difference in score of less than 1.0 between the magnetic bead extraction and vehicle control extraction was considered to indicate that the magnetic bead implants met the requirements of the intracutaneous reactivity test. This intracutaneous irritation testing was conducted in compliance with ISO 10993-10: 2010, Standard, Biological Evaluation of Medical Devices, Part 10: Tests for Irritation and Skin Sensitization.

## Results

### Comfort

The percent of the total stride time spent in stance during treadmill running was evaluated before and after surgery for the leg that received implants (the right leg) in three turkeys. We did not observe a significant surgery main effect (*p* = 0.234, 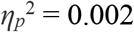) or a significant speed-surgery interaction effect (*p* = 0.492, 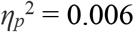) on the percentage of right leg stride time in stance (Figure 5). Out of 600 total strides analyzed (*n* = 20 consecutive strides for three birds and five speeds at each of pre- versus post-surgery biomechanical collections), 5 strides were replaced due to irregular movement (i.e., greater than 3 standard deviations from the mean). Linear mixed model analysis revealed a significant main effect of speed (*p* < 0.001, 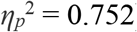) on the percentage of right leg stride time in stance. The percentage of right leg stride time in stance significantly decreased with increasing speed (*p*-values < 0.001).

**Figure 5:**
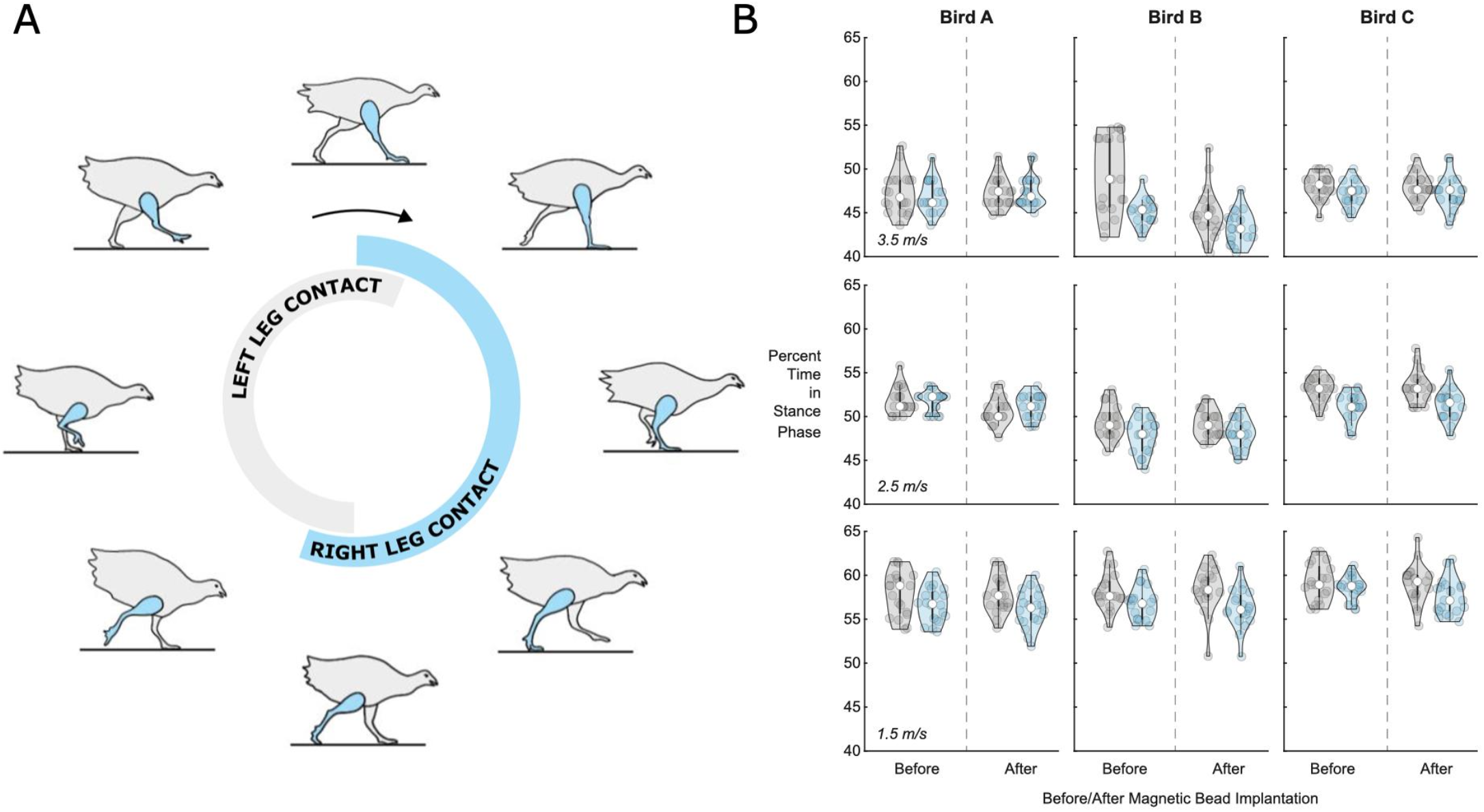
Percentage of Stride Time in Stance Phase Before and After Surgery. **(A)** Turkey gait cycle showing the portion of gait when the leg is in contact with the ground (the stance phase) during running. The contact time for the right leg, which received the implants, is shown in blue, while the contact time for the left leg is shown in gray, for reference. **(B)** Violin plots for left leg (gray, for reference) and right leg (blue) percentage of stride time in stance phase (n=20 strides) for each bird (A, B, and C) at 1.5 m/s (bottom), 2.5 m/s (middle), and 3.5 m/s (top). See supplementary file StancePercent.xlsx for the full dataset in table form.

### Migration

To analyze the effect of distance on migration, we implanted magnetic beads into a total of 16 turkey muscles across seven turkeys. The results of this migration study are shown in Figure 6. Magnetic bead pairs were implanted at various distances apart and their separation distances were determined via CT scan both immediately following implantation and after eight months post-implantation. Full migration occurred in three implants at and below 2.01 cm, but did not occur in one implant at 1.83 cm or in any implants at or above 2.12 cm. As indicated in the plot, one of the muscles was implanted with three magnets in line with one another, for a separate investigation. See Supplementary Table 1 for a numerical presentation of the initial and final separation distances.

**Figure 6:**
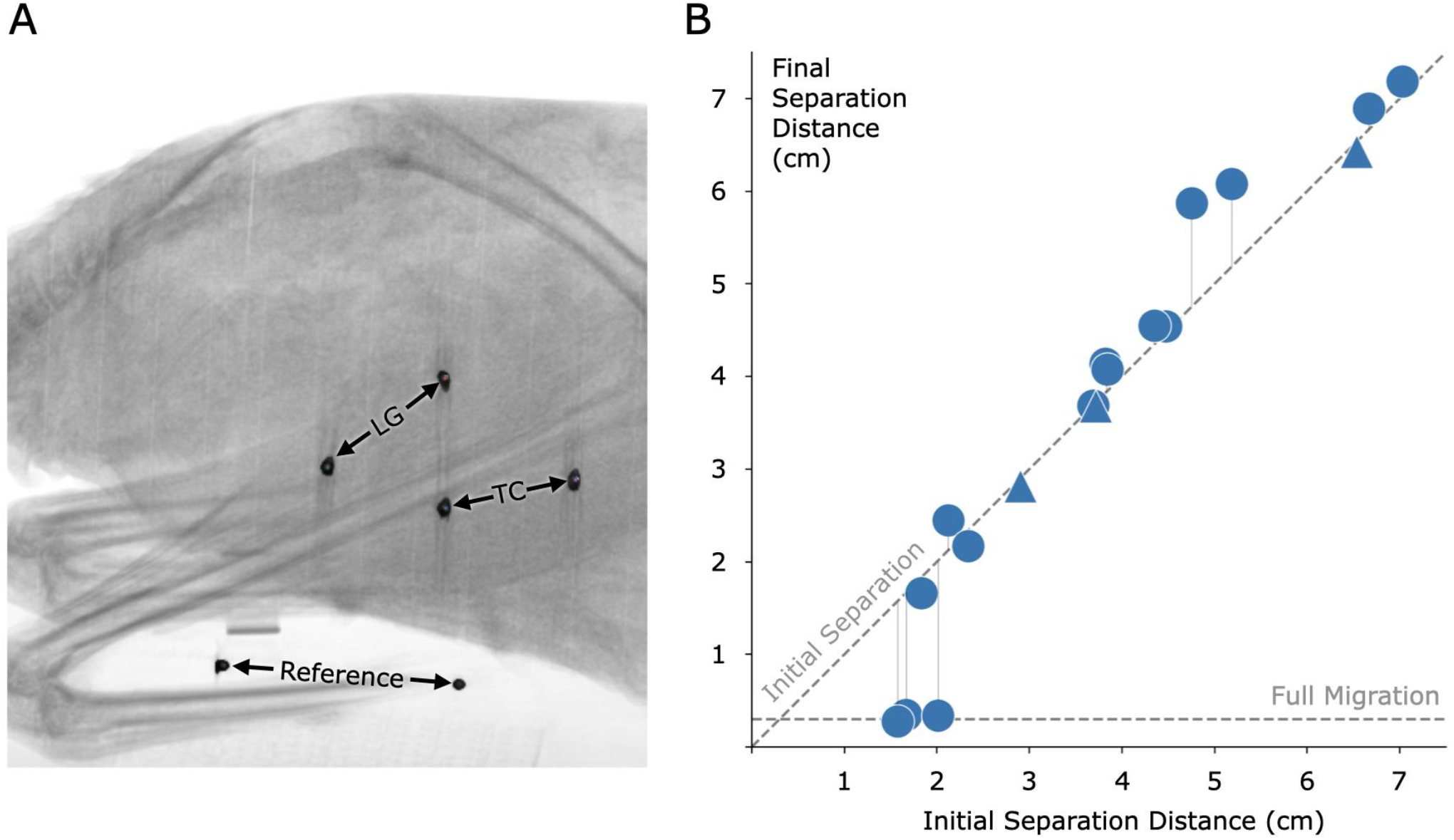
Magnetic Bead Pair Stability Against Migration in Muscle. **(A)** Computed tomography image of one of the turkeys with the implanted magnetic beads. In this case, magnets were implanted in the lateral gastrocnemius (LG) and tibialis cranialis (TC) muscles. A consistent reference, seen at the bottom of the figure, was included in all scans to ensure repeatable distance measurements. **(B)** Effect of initial implant separation on migration of the magnetic beads, seen by comparing the vertical position of each data point to the diagonal dashed line. The horizontal dashed line, labeled Full Migration, indicates the distance between bead centers when they are touching (3 mm center to center). The final magnetic bead pair separation is the separation at eight months post-implantation. We observed a lack of full migration for all implant pairs at or above an initial separation distance of 2.12 cm. Triangles indicate pairwise distances between three magnets that were placed approximately in line within a single muscle. Refer to Supplementary Table 1 for this data in a numerical representation.

### Biocompatibility

For the turkey portion of the biocompatibility analysis, no inflammation was observed around any of the implant sites (see Figure 7 for a representative histological cross section).

**Figure 7:**
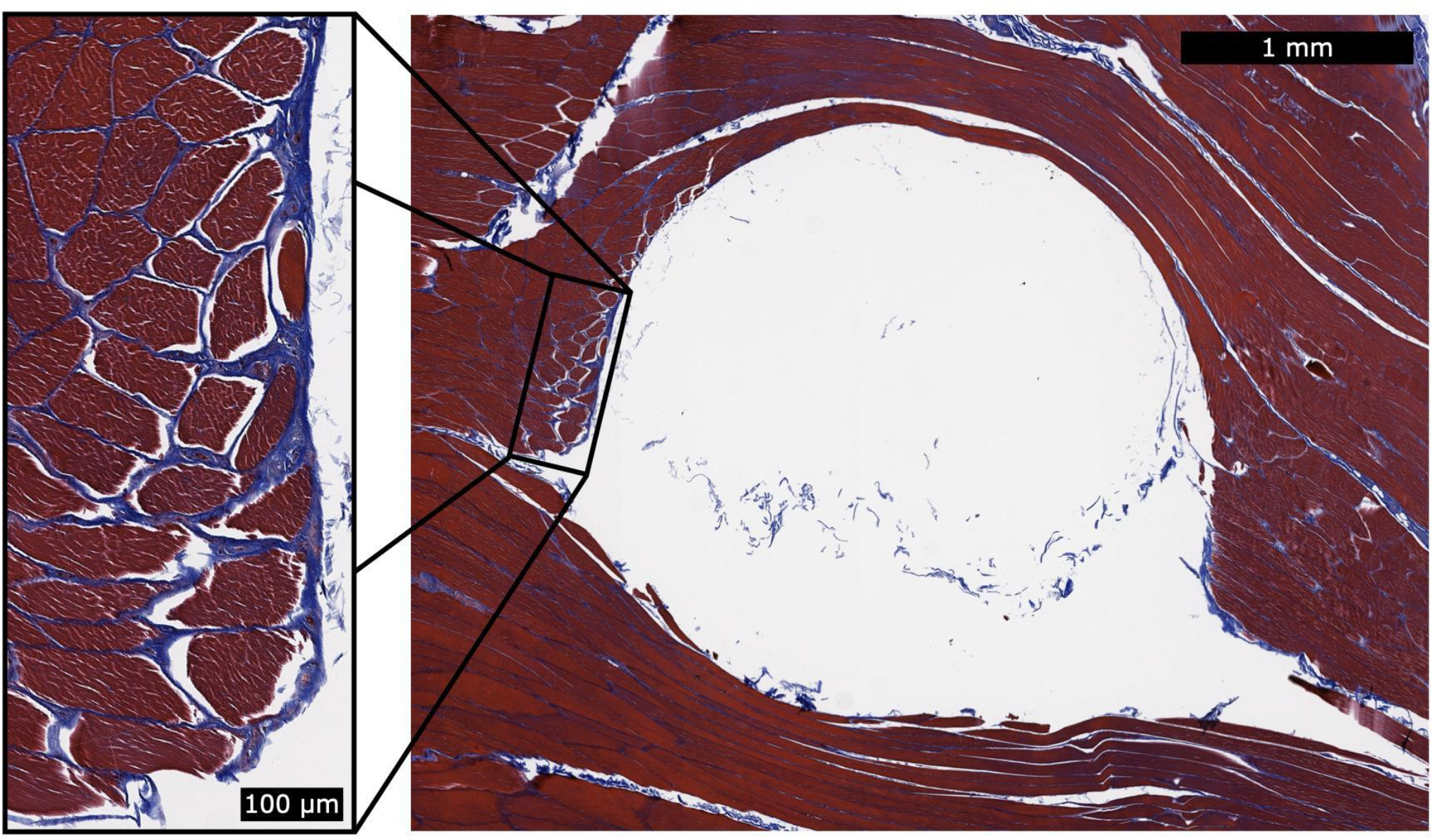
Histology for a Single Magnetic Bead Implant. This representative histology sample shows a cross section of the implantation site after removal of the spherical magnetic bead implant. We applied Masson’s Trichrome staining to highlight the fibrous tissue in blue. During explantation, some of the fibrous capsule remained attached to the implant and thus was generally unobservable in the histological analysis. However, a thin fibrous wall remained (see magnified inset), and the fibrous tissue had integrated into the muscle fibers. This observation suggests that the fibrous tissue may contribute to holding the implant in position relative to the surrounding muscle tissue. No inflammation was observed surrounding the implantation sites.

The GLP testing results demonstrated via extensive biocompatibility testing that the implants were non-irritant, non-cytotoxic, non-allergenic, and non-irritating (see Table 1).

**Table 1:** Biocompatibility Testing Results. The following table lists the tests that were performed on the magnetic bead implants under GLP, along with the ISO standards that were followed and the corresponding test results.

| Test | ISO Standard | Result |
| --- | --- | --- |
| Implantation - 2 wk | 10993-6:2016 | 0.0 (non-irritant) |
| Implantation - 26 wk | 10993-6:2016 | 0.0 (non-irritant) |
| Cytotoxicity | 10993-5:2009 | 0 (non-cytotoxic) |
| Sensitization | 10993-10:2010 | 0 (non-allergenic) |
| Irritation | 10993-10:2010 | 0.0 (non-irritating) |

### Biocompatibility: Intramuscular Implantation Testing

The two-week and twenty-six-week intramuscular implantation tests in rabbits each resulted in an irritant ranking score of 0.0 (rounded up from negative) for the magnetic bead implants, indicating that the fully manufactured magnetic beads in this study are non-irritants as compared to the control articles.

No abnormal clinical signs were noted, and no difference was noted macroscopically between the test and control sites in any of the rabbits. All rabbits survived to the scheduled study endpoint.

### Biocompatibility: Cytotoxicity Testing

The cytotoxicity test resulted in a score of 0 for the magnetic bead extraction at 24, 48 and 72 hours, indicating that the magnetic bead implants of this study are non-cytotoxic under the conditions of the cytotoxicity test. The negative and cell controls also received scores of 0 (no reactivity, discrete intracytoplasmic granules, no cell lysis, and no reduction of cell growth), while the positive control received a score of 4 (severe reactivity and complete or nearly-complete destruction of the cell layers), indicating that the test was functioning normally.

### Biocompatibility: Sensitization Testing

The sensitization test resulted in a score of 0 for dermal observations of all exposure sites for both the normal saline and sesame oil extractions, indicating that the magnetic bead implants did not elicit a sensitization response, and thus that the implants are non-allergenic under the conditions of the test. No abnormal clinical signs were noted, and all animals survived to the scheduled study endpoint.

### Biocompatibility: Intracutaneous Irritation Testing

The intracutaneous irritation test resulted in a comparative score of 0.0 for erythema and edema against the vehicle control for both the normal saline and sesame oil extractions, indicating that the magnetic beads met the requirements of the intracutaneous reactivity test, and thus that the implants are non-irritating under the conditions of the test. No abnormal clinical signs were noted, and no dermal reactions were observed at the test or control sites in any of the rabbits at any of the observation points. All animals survived to the scheduled study endpoint.

## Discussion

In this study, we investigate the clinical viability of magnetic bead implants in muscle. Using a specialized device to insert magnetic beads into muscle in avian and lagomorph models, we collect data to assess bead migration, gait metrics, and bead biocompatibility. We find that implanted-leg stance time is preserved from pre-to post-implantation of the magnetic beads in muscle, migration does not occur when the magnets are implanted a sufficient distance from one another, and no inflammation occurs in response to the magnetic bead implantation. In addition, GLP-compliant testing confirms that the implants are non-irritant, non-cytotoxic, non-allergenic, and non-irritating. These results suggest that MM is a viable approach to muscle tissue length tracking in humans.

### Comfort

In support of our hypothesis, we found no evidence of magnetic bead implant discomfort from measures of pre- and post-implant gait. Following a series of training sessions pre- and post-implantation, we did not observe an effect of surgery on the percentage of right leg stride time in stance across a range of walking and running speeds. The absence of any sign of antalgic gait in a relatively small animal three weeks following implantation supports the idea that the implant would not cause significant discomfort in humans, where the implant would represent a much smaller fraction of muscle volume.

### Migration

In previous work, no magnets migrated toward one another from an initial separation distance of 2.15 cm or above (Taylor et al. 2021). Due to the now increased magnetization strength of the magnetic implants (N48 versus N35, or approximately 1.4 T versus 1.183 T), we hypothesized that no implants at this increased strength would migrate from an initial separation of 2.34 cm or greater, a predicted increase of 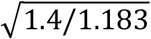, or about 9% (using the migration model of Supplementary Note 1, Taylor et al. 2021). In this investigation, no implants were found to migrate from this distance, but also no implants were found to migrate from the even closer distance of 2.12 cm. Having observed in previous work that one pair of magnets only partially migrated from a starting location of 1.67 cm, we further hypothesized that we may see magnets not fully migrate from an initial separation distance as low as 1.82 cm. In support of this hypothesis, in this study we observed a magnetic bead pair that did not migrate from a starting separation distance of 1.83 cm, even though another bead pair did migrate from 2.01 cm. These results highlight the lack of a precise cutoff for a migration distance threshold, due to the inherent limitations of solely relying on initial separation distance to the exclusion of all other factors.

The migration model used here to predict the migration cutoffs may help in understanding one aspect of migration but is insufficient on its own to fully capture what is happening within the body with magnetic bead implant pairs. While in this study we employed the same material type for coating as in previous work, our updated coating, cleaning, and insertion processes resulted in no observed inflammation, which may have reduced the likelihood of migration. Variations in the magnetization strength of each implant due to magnetic bead size tolerances and magnetic material alignment may also be a cause for both in-study and cross-study variations, as could small differences in the surgical procedure (e.g., the position of the insertion path). Finally, our model holds as a fundamental assumption that the maximum contraction ratio is equivalent between any two beads, regardless of implant location, suggesting that the minimum distance between the beads occurs during full muscle contraction and can be predicted from the distance between the beads at rest. However, muscle contraction is not spatially uniform (for instance, widening as it shortens), which weakens this assumption when the beads are not placed in precisely the same locations within the muscle. Further, this assumption is also weakened by the possibility of subject-to-subject variability in contraction ratios. Importantly, a highly conservative estimate of the migration threshold should be used to account for the many biological factors of the implants and implantation process. As such, we maintain the importance of an initial 3 cm separation distance at the time of implantation for these 3-mm-diameter spherical magnetic beads to ensure stability against migration.

It is critical that these 3-mm-diameter magnetic bead implants be verified to be at least 3 cm from *all* other implants for *all possible joint configurations*, including distinct muscle contraction states resulting in all possible joint configurations. We wish to also underscore the importance of implanting at, or beyond, the minimum distance threshold even when only a single magnetic bead is implanted per muscle, as has been proposed in previous work (Tarantino et al. 2017). It is critical that this verification be included in the surgical implantation protocol design for implantation in all future human studies.

### Biocompatibility

In a previous study, we observed the histological results of implants coated with a proof-of-concept coating and insertion technique. In this investigation, we developed and employed a hospital-ready insertion technique to implant magnetic beads with standard medical coatings of gold and Parylene C. With the improved techniques and protocols of this study, we did not observe inflammation caused by the implants. We contracted a GLP laboratory to formally investigate the implants for biocompatibility using this finalized clinical-grade insertion device and implants, and the implants were shown to be non-irritant, non-cytotoxic, non-allergenic, and non-irritating. These results support the clinical biocompatibility of these implants for human use.

### Limitations

These magnetic bead implants have not yet been subjected to MRI Safety Testing, an important factor in their potential use in humans, as these implants may limit the ability of a patient to get an MRI image.

Until MRI Safety Testing is completed for these devices, a patient would need to be instructed to not get an MRI. Though there are cases in which an MRI is allowed when a patient has a magnetic implant, the imaging conditions are highly specific to the particular geometry, strength, coercivity, location, and setup of the permanent magnets, making a study of MRI compatibility necessary (Edmonson et al. 2018).

This technique does not replace, and should not be seen as a potential replacement for, RSA, which is a standard technique for monitoring migration of implants such as knee replacements (Kärrholm 1989). RSA is an excellent technique for monitoring rigid bodies because it allows many small beads to be implanted at once and at any depth, while magnetic beads must be separated by a minimum distance and have a limited implantation depth. MM is designed for use in situations requiring high-accuracy real-time tissue length tracking in relatively superficial muscles.

Magnetic bead tracking is depth-limited due to the signal-to-noise ratio of the sensors and the nature of the magnetic dipole field, which falls off with the inverse cube of distance. With current sensing technology, magnet tracking is optimal for use at a depth between no less than 6 mm (see Table 5.1 of (Taylor 2020) for the recalibration point when tracking a 1.4 T magnet) and no greater than 33 mm (see Supplementary Figure 1).

In this work, we evaluated 3-mm-diameter spherical magnetic beads coated in Parylene C. Beads of different sizes, geometries, and coatings would need to be further investigated for comfort, migration, and biocompatibility before clinical use.

### Applications

The use of magnetic beads in human-machine interfacing enables additional strategies for controlling external devices and monitoring tissue states. For instance, the use of magnetic bead pairs to track muscle tissue length via MM could allow a robotic prosthesis to be controlled using muscle tissue lengths in paired agonist and antagonist muscles, providing position and impedance control (see Figures 8A and 8B). Similarly, muscle tissue lengths could be used to control an exoskeleton to provide restoration or augmentation of weak or healthy muscle movement (see Figure 8C). Magnetic bead implants could also enable closed-loop artificial muscle stimulation, providing feedback about the muscle’s length during stimulation to allow high-fidelity control over the muscle (see Figure 8D).

**Figure 8:**
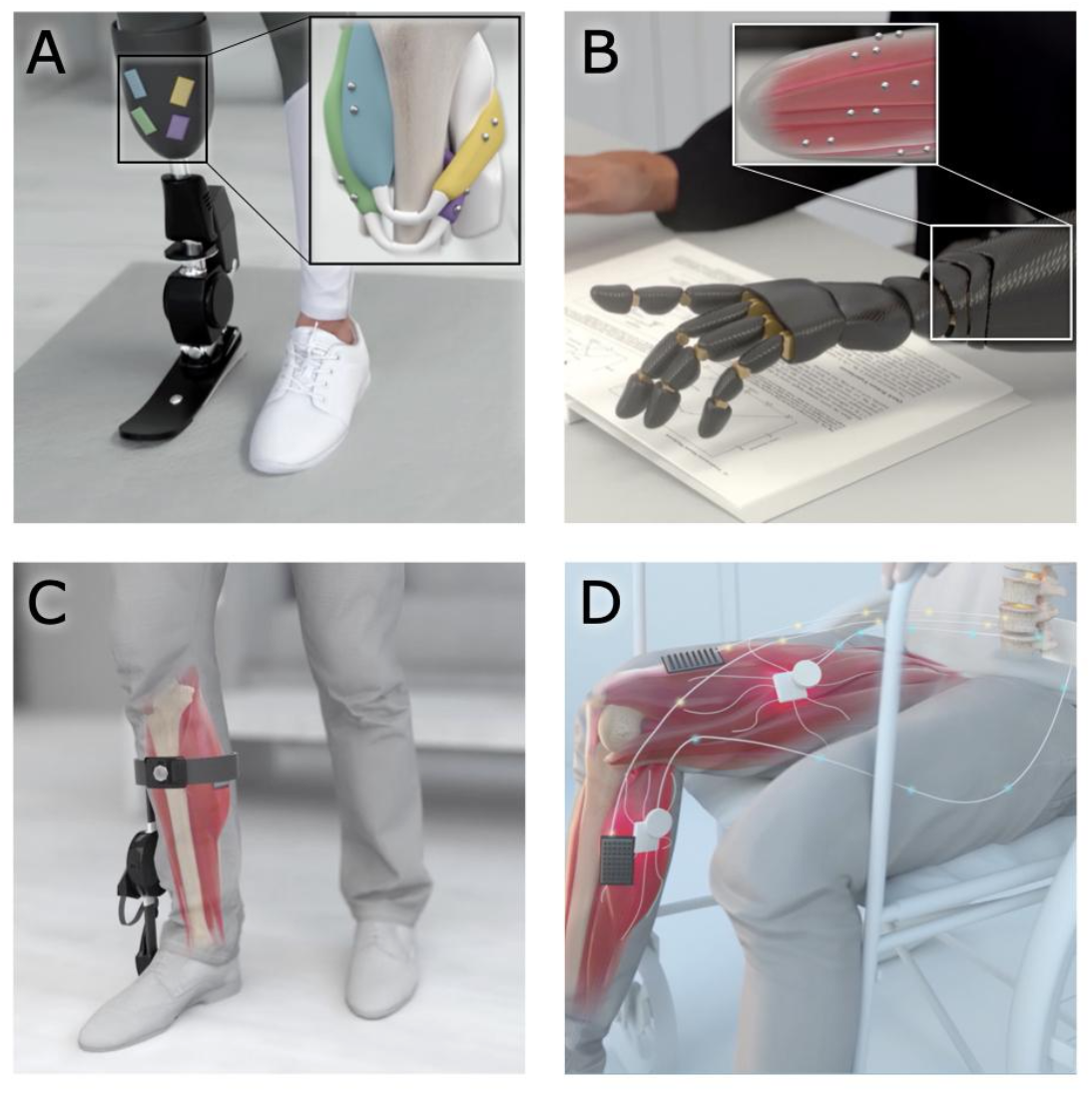
Applications of Magnetomicrometry. When used to track muscle tissue lengths via magnetomicrometry, magnetic bead implants could enable real-time control in human-machine interfaces. **(A, B)** Magnetic beads implanted into residual muscles could be used to control a prosthetic limb device. **(C)** When implanted in a weakened muscle, the magnetic beads could provide control over an exoskeleton for restoration or augmentation of joint torque. **(D)** Magnetic beads in paralyzed muscles could enable closed-loop artificial muscle stimulation for control of muscle length or force.

While the more urgent applications of muscle tracking address the restoration of natural human abilities, muscle tissue length tracking could enable various human-machine interface tasks, such as the operation of remote machines (e.g., unmanned aerial vehicles or telepresence robots), operation of local machines (e.g., factory or farm equipment), and control of oneself or one’s tools in an augmented or virtual reality environment. Muscle length control could also enable improved navigation in a weightless environment, such as in control of underwater thrusters or air thrust rotors. Specifically, muscle tissue length tracking via magnetic beads is unaffected by water, in comparison with surface electromyography, where the leads need to be fully waterproofed to work reliably when submerged (Rainoldi et al. 2004). In all these cases, the use of muscle tissue length to control an external device would provide an intuitive and powerful means of interaction, as humans are already proficient at using muscle length to control their own movements.

Magnetic bead implants may also be used for delivering proprioceptive feedback. Illusory kinesthetic feedback via muscle tissue vibration has been well tested via vibramotors on the skin surface, and works by activating muscle spindles to deliver the sensation of illusory joint positions and movements (Marasco et al. 2018). However, as currently implemented, vibration through the skin may create undesirable cutaneous sensations, distracting from the desired proprioceptive signal. Implanted magnetic beads provide an opportunity to vibrate a muscle from the inside using electromagnetic actuation. The ability to selectively vibrate individual magnets was recently demonstrated in benchtop tests of a “myokinetic stimulation interface,” including the ability to modulate the direction of the vibration (Montero et al. 2021). For translation, this strategy will need to balance weight and power requirements, manage simultaneous tracking and stimulation, and compensate for vibration of the surrounding electromagnets. Once these issues are addressed, these passive intramuscular magnetic beads could be employed as bidirectional human-machine interfaces.

When implanted next to cutaneous receptors, such as in fingertips, magnetic implants can confer cutaneous sensing of low-frequency magnetic fields (Hameed et al. 2010). Seeking this additional magnetic sense, the biohacking community has been implanting magnets since the late 1990s or early 2000s (Doerksen 2018), though these implantations are performed as a do-it-yourself operation (Yetisen 2018) and often without anesthesia (Brickley 2019). While we advise against the self-implantation of non-clinical-grade magnetic beads, this history of magnetic bead implantation suggests an innate human desire to be augmented and a simplicity to the use of passive magnetic beads as a human-machine interface.

### Summary

In this work, we manufacture clinical-grade magnetic bead implants and develop a clinical-grade implantation strategy, and we verify implant comfort, stability against migration, and biocompatibility. Our results demonstrate that when implanted as discussed here, these magnetic beads are viable for use in human muscle.

## Supporting information

StancePercent

## Author Contributions

CT developed the coating strategy and implant manufacturing process, co-designed the custom magnetic bead inserter, coordinated the GLP biocompatibility testing, oversaw experimental design, contributed to surgical procedure design, assisted in surgeries, oversaw the performance of histology, aided in computed tomography data collection, performed migration data analyses, maintained experimental documentation, and led the manuscript preparation. WC contributed to experimental design, assisted in surgeries, performed ultrasound measurements, collected gait symmetry data, performed gait symmetry statistical analyses, performed computed tomography data collection, and contributed to manuscript preparation. EC assisted in collecting and processing gait symmetry data and contributed to manuscript preparation. SY collected and analyzed the magnetomicrometry depth dataset and performed magnetomicrometry measurements during surgeries. MC contributed to surgical procedure design, assisted in surgeries, and contributed to manuscript preparation. SL oversaw the statistical design and statistical analysis of the gait symmetry experiments. RB performed histological analyses and contributed to manuscript preparation. TR assisted with general study management, contributed to experimental design, assisted in surgeries, aided in performing experiments, and contributed to manuscript preparation. HH conceived of the study, oversaw project funding, assisted with general study management, contributed to experimental design, and aided in manuscript preparation.

## Acknowledgments

The authors thank, non-exhaustively, Alexander Best, Mckay Bruning, Andreas Burger, Ziel Camara, Tyler Clites, Steven Charlebois, Charlene Condon, Kathy Cormier, Bruce Deffenbaugh, Michelle Dietzel, Asami Ehlert, MacKenzie Ess, Rachel Fleming, Lisa Freed, Robert Gnos, Deborah Grayeski, Alan Grodzinsky, Guillermo Herrera-Arcos, Crystal Jones, Kylie Kelley, Aimee Liu, Richard Marsh, Jesse Mendel, Richard Molin, Chad Munro, Allison O’Konek, Jarrod Petersen, Mitchel Resnick, Lindsey Reynolds, Andy Robinson, Emily Rogers, Amy Rutter, Jessica Sohner, Shriya Srinivasan, Erika Tavares, and Beni Winet for their helpful advice, suggestions, feedback, and support. Inclusion in this list of acknowledgments does not indicate endorsement of this work.

## Data Availability

The original contributions presented in the study are included in the article and supplementary materials. Further inquiries can be directed to the corresponding author.

## Supplementary Material

**Supplementary Figure 1:**
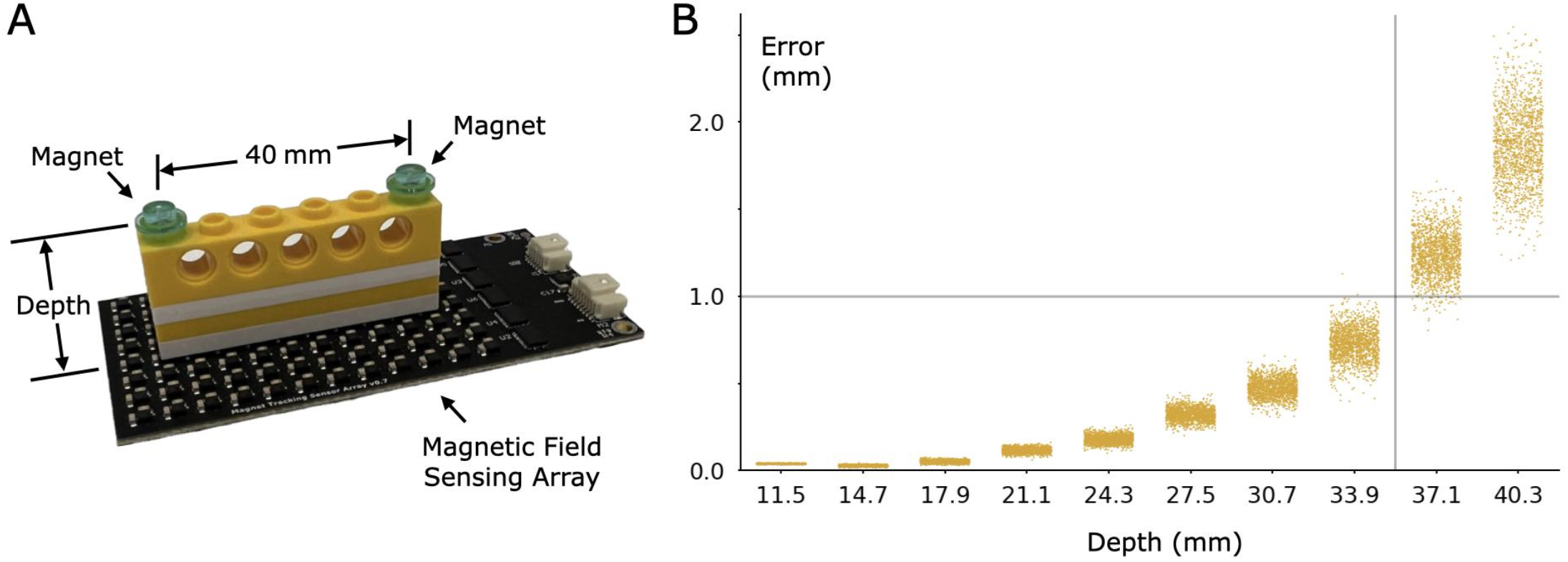
Magnetomicrometry Sensing Error vs Implantation Depth of Two Implantable Magnetic Beads. **(A)** We affixed two magnetic beads (manufactured as described in the methods) into 1×1 LEGO round plates, and we attached these two round plates 40 mm apart to a 1×6 LEGO technic brick. We then centered this LEGO brick over a 96-element (LIS3MDL) magnetic field sensing array parallel with the long axis of the array. We varied the depth in 3.2 mm increments from the initial depth of 11.5 mm by adding 1×6 LEGO plates. This centering presents a best-case scenario, suggesting maximum depth limits that could be achieved with proper placement of magnetic field sensors. **(B)** Seaborn strip plots show the error in mm (vertical axis) of the magnetomicrometry signal for each depth. The horizontal axis shows each depth categorically. Note that the magnetomicrometry best-case accuracy with this sensing array and these magnets is sub-millimeter to a depth of about 33 mm.

**Supplementary Table 1:** Initial and Final Separation Distances Between Implanted Magnets. The following table lists the various magnetic bead separation distances sorted in ascending order by initial separation distance. We implanted magnetic bead pairs into the lateral gastrocnemius (LG) and tibialis cranialis (TC) of the right (R) legs of turkeys D-F. We implanted a magnetic bead pair into the right LG and three magnetic beads into the left (L) LG of turkey H. For the left LG of turkey H, we have listed all three pairwise initial and final separation distances between the three implanted magnetic beads. We implanted magnetic bead pairs into the femoralis tibialis (FT) and the LG of both legs of turkey I and in the left leg of turkey J.

| Separation (mm) |  | ID/Leg/Muscle |
| --- | --- | --- |
| Initial | Final |  |
| 15.7 | 2.8 | I/L/LG |
| 16.6 | 3.5 | J/L/LG |
| 18.3 | 16.6 | J/L/FT |
| 20.1 | 3.4 | I/R/FT |
| 21.2 | 24.5 | I/L/FT |
| 23.4 | 21.7 | I/R/LG |
| 29.0 | 28.2 | H/L/LG (2 to 3) |
| 36.9 | 36.9 | G/R/TC |
| 37.2 | 36.8 | H/L/LG (1 to 2) |
| 38.2 | 41.4 | D/R/LG |
| 38.4 | 40.8 | F/R/TC |
| 43.5 | 45.5 | G/R/LG |
| 44.7 | 45.4 | F/R/LG |
| 47.5 | 58.7 | H/R/LG |
| 51.9 | 60.8 | D/R/TC |
| 65.3 | 64.3 | H/L/LG (1 to 3) |
| 66.6 | 69.0 | E/R/LG |
| 70.2 | 71.8 | E/R/TC |

